# Reconstructing the hyphosphere using a hyphal release-capture soil microcosm

**DOI:** 10.64898/2026.05.26.727979

**Authors:** Gayan Abeysinghe, Elek Nagy, Tanya Wagner, Shravan Parunandi, Joshua Santos, Muthukumar Bagavathiannan, Sanjay Antony-Babu

## Abstract

Fungal hyphae form spatially confined interfaces in soil that mediate close associations with bacteria, collectively referred to as the hyphosphere. Despite its recognized ecological importance, experimental access to hyphosphere-associated microbial communities under realistic soil and plant-associated conditions has remained limited. Here, we present a soil-mimetic microcosm that enables controlled reconstruction and recovery of hyphosphere bacterial communities embedded within plant-associated soil. The system integrates field-derived soil, a native soil microbial inoculum, living cotton seedlings, and a spatially constrained fungal inoculum housed within sterile cell-strainer assemblies, permitting hyphal extension into soil while preserving a recoverable fungal-soil boundary. Using the soil-borne plant pathogen *Fusarium oxysporum* f. sp. *vasinfectum* as a model filamentous fungus, we show that the microcosm enables reproducible recovery of hypha-associated soil microaggregates containing physically attached bacterial cells. Full-length 16S rRNA profiling revealed pronounced reductions in bacterial richness and evenness in hyphosphere samples relative to bulk and rhizosphere soils, consistent with recruitment of a restricted subset of the surrounding microbiota. Ordination analyses demonstrated clear compositional separation between soil and hyphosphere compartments, with convergence of hypha-associated communities across bulk and rhizosphere contexts. Phylogenetic turnover analyses indicated phylogenetic structuring, whereas taxonomic analyses identified a conserved set of bacterial genera consistently associated with hyphae alongside compartment-specific taxa influenced by soil and plant context. Together, these findings establish the novel hyphal release-and-capture microcosm as a reproducible, ecologically grounded platform for studying hyphosphere-associated bacterial communities in plant-associated soils.

## Introduction

Soils host extraordinarily complex microbial assemblages structured across microscale physicochemical gradients that regulate decomposition, nutrient fluxes, and plant health (Fierer, 2017; Delgado-Baquerizo et al., 2020). Within this heterogeneous matrix, filamentous fungi exert disproportionate influence by creating linear, semi-continuous microhabitats as their hyphae traverse soil pores, translocate water and solutes, and reorganize local microbial distributions (de Boer et al., 2005; Warmink & van Elsas, 2009; Worrich et al., 2016). The narrow fungal interface termed the hyphosphere is now recognized as a distinct microbial niche shaped by hyphal physicochemistry, spatial confinement, and targeted metabolic exudation (Pugh and Sewell, 1958; Toljander et al., 2006; Zhang et al., 2014; Nguyen, 2023; Antony-Babu et al., 2025). In contrast to the rhizosphere, where plant-derived carbon dominates, the hyphosphere forms around discrete hyphal tips and cords, producing unique gradients of organic acids, sugars, amino acids, peptides, siderophores, and secondary metabolites that selectively enrich for compatible bacterial partners (Deveau et al., 2018; Jin et al., 2024).

Fungal-bacterial associations in the hyphosphere are non-random, where they have been well demonstrated to show strain specific selectivity (Rudnick et al., 2015; Thomas and Antony-Babu, 2024). Fungi shape and assemble their hyphosphere microbiota through selective exudation, surface chemistry, and spatial filtering. The recruited bacteria may enhance fungal nutrient acquisition, provide vitamins and siderophores, modulate growth via signaling interference, or suppress fungal expansion through antibiosis or mycophagy (Leveau & Preston, 2008; Frey-Klett et al., 2011; Uehling et al., 2019; Abeysinghe et al., 2020; Uehling et al., 2019). These interactions reverberate through soil-plant systems by altering competition, resource flows, and disease outcomes.

Increasingly, plant pathogens such as members of the *Fusarium oxysporum* species complex are understood not as autonomous agents but as participants within multipartite pathobiomes, where associated microbes influence infection success, host range, and virulence expression (Edel-Hermann & Lecomte, 2019; Berg et al., 2020; Thomas et al., 2025). Among these, *Fusarium oxysporum* f. sp. *vasinfectum* race 4 (FOV4) has emerged as a particularly destructive pathogen of cotton, causing severe yield losses across multiple production regions. Our recent study demonstrated microbial communities associated with its hyphae while within the cotton plant tissues (Antony-Babu et al., 2025), raising the question of which bacteria associate with its hyphae during soil colonization - the stage at which the pathogen first encounters the resident soil microbiota.

Despite their ecological and pathological importance, hyphosphere bacteria remain experimentally challenging to study in ecologically relevant contexts. Current methodological approaches address specific aspects of fungal-bacterial interactions but fall short of capturing hyphosphere assembly in plant associated soils. Bulk soil or rhizosphere sampling obscures hypha-specific recruitment patterns, while culture-based approaches disproportionately recover fast-growing generalists. Microfluidic or fungal highway column systems effectively demonstrate bacterial dispersal along fungal networks, but do not allow targeted study of specific fungal species, instead relying on indigenous fungi that colonize the experimental system (Kohlmeier et al., 2005; Junier et al., 2021; Kelliher et al., 2025). Conversely, “hyphal baiting” strategies enable controlled introduction of specific fungal species into soil matrices (Rudnick et al., 2015; Ghodsalavi et al., 2017), but most implementations study fungal-bacterial interactions in plant-free microcosms a significant limitation given that plant root exudates, rhizosphere chemistry, and soil conditioning by living roots may fundamentally alter hyphosphere community assembly. While some studies have incorporated plant systems (Sun et al., 2021), the hyphal baiting typically occurs in separate, plant-free compartments that do not capture the integrated plant-soil-fungal environment where hyphosphere assembly naturally occurs. This challenge is compounded for regulated plant pathogens such as FOV4, where biosafety restrictions preclude deliberate field inoculation, making controlled microcosm systems the only viable platform for studying hyphosphere assembly under realistic soil conditions. This methodological gap is particularly problematic for understanding pathobiome dynamics, where plant-pathogen-microbiome interactions operate as integrated systems. Consequently, our understanding of how hyphosphere bacterial communities establish and influence fungal ecology in living plant-soil systems remains limited.

Here, we introduce a release-and-capture approach that permits fungal hyphae to extend into soil and enables subsequent recovery of hyphae together with their associated hyphosphere bacterial communities. This method includes a soil-mimetic microcosm explicitly designed to reconstruct hyphosphere assembly under plant-associated soil conditions. This system integrates three essential elements: (i) field-derived soils inoculated with a soil-tea microbial community, to recreate microbial diversity while permitting controlled experimental replication; (ii) cotton seedlings to provide plant-conditioned nutrient profiles, exudates, and microbial structure, as plant roots significantly shape bacterial community composition via root exudation and habitat filtering (Haichar et al., 2008; Guyonnet et al., 2018) and (iii) a spatially delimited fungal inoculum housed within sterile cell strainers that constrain hyphal egress and define a discrete, recoverable fungal-soil boundary. This configuration establishes a controlled but ecologically grounded spatial context, enabling fungal hyphae to extend into a naturalistic microhabitat while preventing uncontrolled soil mixing. Critically, the physical barrier provided by the strainer permits precise isolation of hypha-associated microaggregates, providing an experimentally tractable platform for studying hyphosphere assembly under ecologically relevant conditions.

## Materials and Methods

### 2.1 Overview of method

The overall experimental workflow is summarized in Fig. 1. Since our primary goal is to identify bacteria that are recruited to fungal hyphae from the surrounding soil microbiota, the system must separate the contributions of the soil matrix, the microbial community, and the fungal host so that each can be controlled independently. To study hyphosphere bacterial recruitment under controlled but ecologically relevant conditions, we developed a soil-mimetic microcosm that decouples the soil matrix from its native microbial community and reconstitutes them separately.

**Figure 1.**
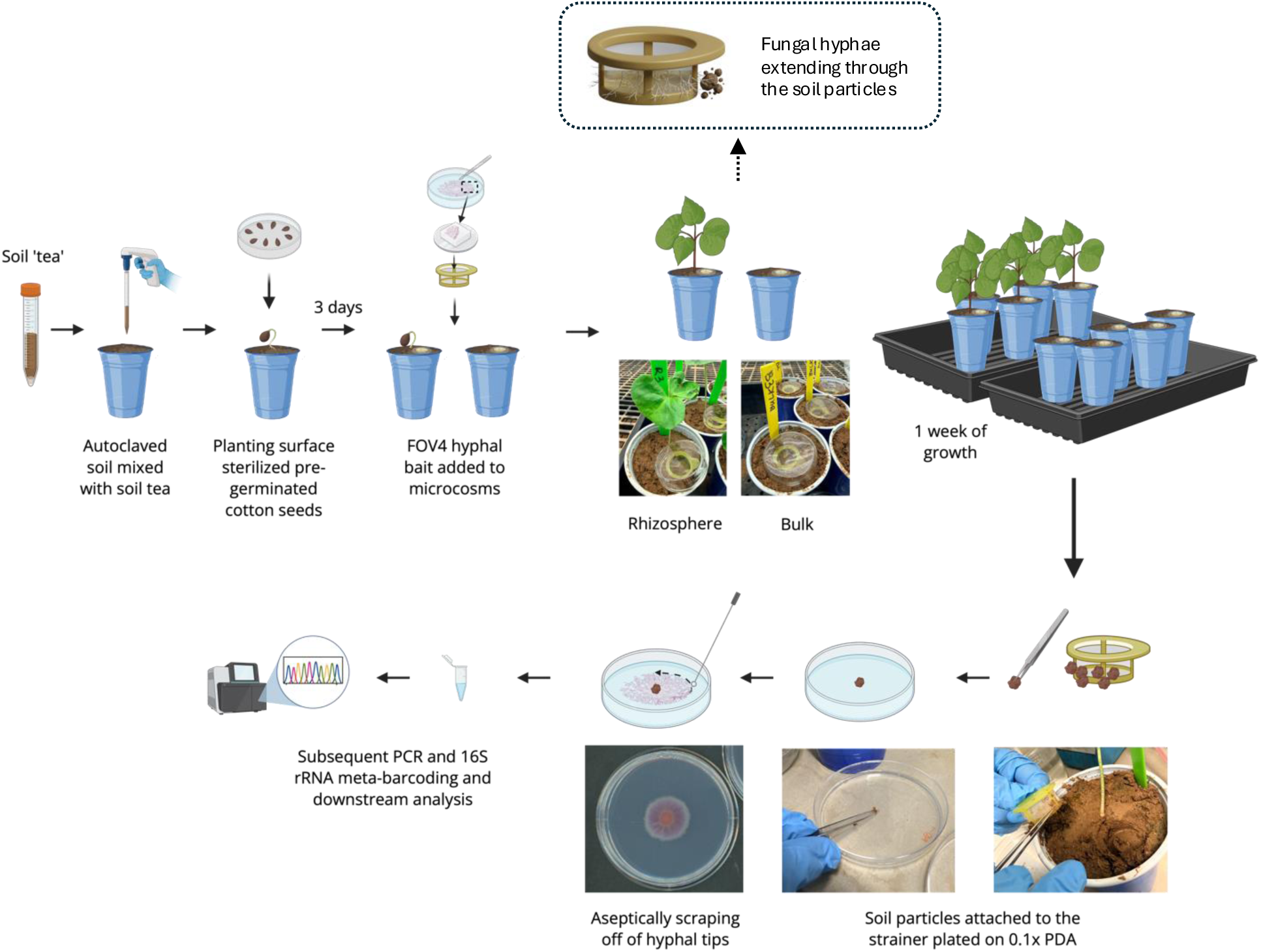
Soil-mimetic microcosm enabling controlled reconstruction and retrieval of hyphosphere-associated bacterial communities. Schematic overview of the experimental workflow used to reconstruct the hyphosphere under plant-associated / non associated soil conditions and recover bacteria recruited to Fusarium oxysporum f. sp. vasinfectum race 4 (FOV4) hyphae. Field soil was homogenized with sterile sand (3:1), inoculated with a soil-tea microbial community, and dispensed into sterile pots. Pre-germinated cotton seedlings were transplanted into designated microcosms and grown for 3 days to establish plant-conditioned soil structure. A sterile 100-µm cell strainer with a sterile coverslip underneath, containing a 3-day-old FOV4 agar plug (0.1× PDA) was inserted either adjacent to seedlings (plant-associated treatment) or centrally in bulk soil (no-plant control). The strainer physically delimited the fungal-soil boundary while permitting controlled hyphal egress into the surrounding substrate. After incubation, strainer assemblies were removed, and hypha-associated soil microaggregates adhering to the rim were transferred onto 0.1× PDA to establish cultures. Actively growing hyphal margins were collected to release the attached bacterial community for culture-dependent isolation and full-length 16S rRNA metabarcoding. Representative images illustrate microcosm configurations and hyphal extension dynamics within the delimited interface.

Field soil was autoclaved to eliminate indigenous microbiota and provide a standardized physicochemical substrate, then re-inoculated with a soil-tea suspension derived from the same field site to introduce a representative but experimentally uniform microbial community across replicates. Pre-germinated cotton seedlings were established in a subset of microcosms to generate plant-conditioned soil. A pathogenic fungal inoculum was introduced via sterile cell strainers that permitted hyphal extension into the surrounding soil while maintaining a recoverable fungal–soil boundary. After incubation, hypha-associated soil microaggregates were recovered for microscopy, culture-dependent isolation, and full-length 16S rRNA gene metabarcoding.

### 2.2 Field soil tea as microbial inocula

Soil samples were collected from the cotton field at the Texas A&M AgriLife Research Field, Brazos Bottom, Texas, USA (30°32′14.0″ N, 96°25′40.1″ W), a site under active cotton cultivation, characterized by clay loam soils. Surface litter and plant residues were removed prior to sampling to avoid contamination from aboveground debris. Soils were obtained from a depth of 5-15 cm at multiple random points (n = 10) to account for spatial heterogeneity and homogenized in sterile bags to generate representative composite samples for downstream analyses. Soils were transported on ice to the laboratory (within 2 hours). To prepare soil-tea inoculum, 6 g of homogenized soil was mixed with 10 mL of sterile 1x phosphate-buffered saline (PBS) supplemented with 30% glycerol to enhance microbial preservation and osmotic stability. The mixture was agitated on a vortex for 5 min at room temperature to detach microorganisms. The suspension (“soil tea”) was used as the inoculum for hyphal baiting microcosms. Aliquots not used on the same day were stored at −80 °C.

### 2.3 Seed surface sterilization and pre-germination

Pima cotton seeds (*Gossypium barbadense* cv. Pima S7) were surface sterilized and pre-germinated prior to microcosm establishment. Surface contaminants were removed by immersing seeds in 1.25% w/v NaOCl for 3 min with gentle inversion, followed by 3 min exposure to 70% (v/v) ethanol. The seeds were then rinsed three times with sterile reverse-osmosis (RO) water to remove residual sterilant. Sterilized seeds were placed on 2% (w/v) Gelrite (five seeds per plate) and incubated in the dark at room temperature for 48 h until radicle emergence. Uniformly germinated seedlings were then transferred into the hyphal-baiting microcosm systems described in Section 2.3.

### 2.4 Microcosm setup

Microcosms were assembled with a 3:1 mixture of pre-autoclaved field soil (121°C for 60 min on 3 consecutive days) and sterile sand. The soil mixtures were dispensed in 450 mL sterile plastic pots. This mixture was inoculated with 5mL of soil tea (Section 2.2) to simulate natural microbial colonization prior to hyphal baiting. Two treatments were established (n = 5 each): (i) bulk soil microcosms without plants; and (ii) planted microcosms containing pre-germinated Pima cotton. For the planted microcosms, pre-germinated Pima cotton seedlings (Section 2.3) were aseptically placed into 3 cm-deep planting holes made with a sterile spatula and were allowed to grow for 3 days. Hyphal baiting assemblies were prepared using sterile 100 µm cell strainer (Falcon 100 µm; CLS352360, Corning, Millipore-Sigma, USA) with a sterile glass coverslip at the base to prevent direct soil contamination. An agar plug (1.5 × 1.5 cm) excised from the colony margin of a 3-day old culture of FOV4 (isolate F4AB; Antony-Babu et al., 2025) grown on 0.1x PDA was placed inside the strainer, positioned with its edge ∼1 mm from the inner rim. The assembly was inserted into a 3 cm deep hole ∼2 cm from the seedling base. The plug was positioned with its edge ∼1 mm from the inner rim of the strainer facing the hypocotyl, enabling controlled hyphal extension through the strainer mesh into the surrounding soil while the mesh maintained a defined, recoverable boundary between the fungal inoculum and soil environment. For bulk-soil (no-plant control) microcosms, the same assembly was inserted centrally, maintaining identical depth and spatial orientation to ensure comparable diffusion and colonization conditions. Each pot received ∼50 mL of sterile 0.25x Hoagland’s solution (Hoagland and Arnon, 1950), providing consistent hydration and maintained in growth chambers, under controlled photoperiod (12.5 hr light/11.5 hr dark cycle) and temperature (23 °C day/18 °C night) for the duration of the experiment.

### 2.5 Hyphosphere culturing

Microcosms were deconstructed aseptically 1 week after the introduction of F4AB baiting assembly to the microcosm. Cell strainer assemblies were removed from the substrate, and soil particles adhering to the outer strainer walls and mesh, bound by hyphae that had extended through the strainer into the surrounding soil, were gently removed using sterile forceps. These hyphal-associated soil aggregates were placed at the center of two 0.1x potato dextrose (PD) agar plates (1.5% w/v agar) and incubated at 28 °C until visible hyphal outgrowth extended beyond the original aggregates. Actively growing hyphal tips were then aseptically transferred using sterile toothpicks into microcentrifuge tubes containing 500 µL of molecular-grade water. Each sample was vortexed for 5 s and pipetted up and down five times to dislodge hyphosphere-associated bacteria and spores, generating a uniform hyphosphere suspension. A 250 µL aliquot of each suspension was mixed with 250 µL of 60% glycerol to prepare long-term glycerol stocks of the hyphosphere community. Fungal identity in representative samples was confirmed as *F. oxysporum* by ITS sequencing. The remaining suspensions were immediately used for DNA extraction and 16S rRNA gene metabarcoding.

### 2.6 Microscopy of hyphosphere-associated bacterial communities

Hyphosphere-associated bacteria were visualized from cultures derived from hypha-bearing soil microaggregates (Section 2.5). Following 48 h outgrowth on 0.1x PDA, actively extending hyphal margins with adherent bacterial cells were imaged directly on agar surface using an Echo Revolution microscope (Echo, USA) in upright configuration. Bright-field images were acquired using a 20x/0.45 NA Plan Fluorite Ph1 objective (OFN22). Images were captured using the native Echo acquisition software, and only linear adjustments to brightness and contrast were applied uniformly across all samples. Scale bars were generated automatically from the calibrated optical magnification.

### 2.7 DNA extraction and long-read metabarcoding sequencing

DNA was extracted from the hyphosphere suspensions (Section 2.5) by heat lysis at 95 °C for 5 minutes with intermittent vortexing (Antony-Babu et al., 2025). The presence of bacterial DNA associated with fungal hyphae was verified by amplifying the 16S rRNA gene using universal bacterial primers 27F (5′-AGA GTT TGA TCM TGG CTC AG-3′) and 1492R (5′-TAC GGT ACC TTG TTA CGA CTT −3′). PCR amplifications were performed using the KAPA 2G HotStart ReadyMix with Dye (KAPA Biosystems, USA) under the following cycling conditions: initial denaturation at 95 °C for 3 min, followed by 30 cycles of 95 °C for 15 s, 53 °C annealing for 15 s, and 72 °C for 1 min, with a final extension at 72 °C for 5 min. Amplicons (∼1400-1600 bp) were visualized on 1% (w/v) TBE agarose gels. 16S rRNA gene amplicons were used to construct long-read libraries using the Oxford Nanopore Rapid Barcoding Kit (SQK-RBK114.24, ONT) following the v14 chemistry workflow. A final pooled library was loaded onto a PromethION R10.4.1 flow cell and sequenced using a PromethION P2 device. Raw POD5 data were basecalled and demultiplexed using Dorado v0.9.1 with the super-accurate (SUP) model appropriate for R10.4.1. Reads were processed using NanoFilt v1.1.0, retaining a Q-score ≥ 10.

### 2.8 Bioinformatics pipeline and data analysis

For comparative analysis, samples were grouped into four categories: bulk soil (BS), bulk soil hyphosphere (BH), rhizosphere soil (RS) and rhizosphere soil hyphosphere (RH). Sample size was n=5 per group (BS, BH, RS, RH; 20 samples total). Taxonomic classification was performed using EMU v3.5.1 against the rrnDB v5.6 and NCBI Targeted Loci databases, retaining taxa with relative abundance ≥0.001% to minimize spurious classifications while maintaining rare biosphere detection. Reads were length filtered to retain sequences between 1200-1800 bp.

All analyses were performed in Python 3.12 using pandas 2.2.3, scipy 1.14.1, scikit-bio v0.6.0, numpy 1.26.4, matplotlib 3.10.0, and seaborn 0.13.2. Alpha diversity (richness, Shannon, Pielou’s evenness) was calculated at genus level, with differences tested using Kruskal-Wallis followed by pairwise Mann-Whitney U tests (α = 0.05). Beta diversity was assessed via Bray-Curtis dissimilarity and visualized by PCoA. Compositional differences among compartments were tested using PERMANOVA (scikit-bio v0.6.0; 999 permutations) for the omnibus four-group test and all pairwise comparisons, with p-values and Benjamini-Hochberg FDR-corrected q-values.

Homogeneity of multivariate dispersion was assessed using PERMDISP on the same Bray-Curtis matrix, with within-group dispersion summarized as sample-to-centroid distances. Hierarchical clustering used UPGMA linkage. Indicator species analysis identified compartment-specific genera (IndVal ≥0.5, p < 0.05) and differential abundance was assessed via log₂ fold change.

Phylogenetic community assembly was evaluated using beta nearest taxon index (βNTI) following Stegen et al. (2013). Reads were clustered at 95% identity using VSEARCH to generate OTU reference. These were aligned using MAFFT and a phylogenetic tree was inferred using FastTree and subsequently pairwise phylogenetic turnover among communities was quantified using abundance weighted βMNTD. βNTI was calculated using 999 null randomizations, with values <-2, −2 to 2 and >2 indicating homogeneous selection, stochastic assembly and variable selection respectively. Phylogenetic turnover was evaluated within compartments and across key compartment transitions, including bulk soil to bulk hyphosphere (BS vs BH) and rhizosphere to rhizosphere hyphosphere (RS vs RH).

## Results

### Microcosm design enables controlled reconstruction of hyphosphere communities

The soil-mimetic microcosm produced a spatially defined hyphal-soil interface that was consistent across both planted and plant-free conditions. In planted-microcosms, the cotton seedlings established root system within 3 days, generating a rhizosphere distinct from the plant-free soils.

Following the one week incubation, the FOV4 hyphae extended through the strainer mesh to extend freely into the surrounding soil, and hypha-associated soil microaggregates adhering to the strainer rim were readily visible and could be aseptically recovered. These microaggregates consistently yielded FOV4 outgrowth accompanied by adherent bacterial cells on low-nutrient agar, indicating that intact hyphal-bacterial associations formed within the soil were preserved during recovery.

Brightfield microscopy confirmed the physical attachment of bacterial cells along hyphal surfaces following plate outgrowth, including localized microcolonies along lateral walls and hyphal tips (Fig. 2A). Together, these observations validate that the microcosm supports *in situ* reconstruction of the hyphosphere assemblage and enables downstream recovery of hypha-associated bacterial communities.

**Figure 2.**
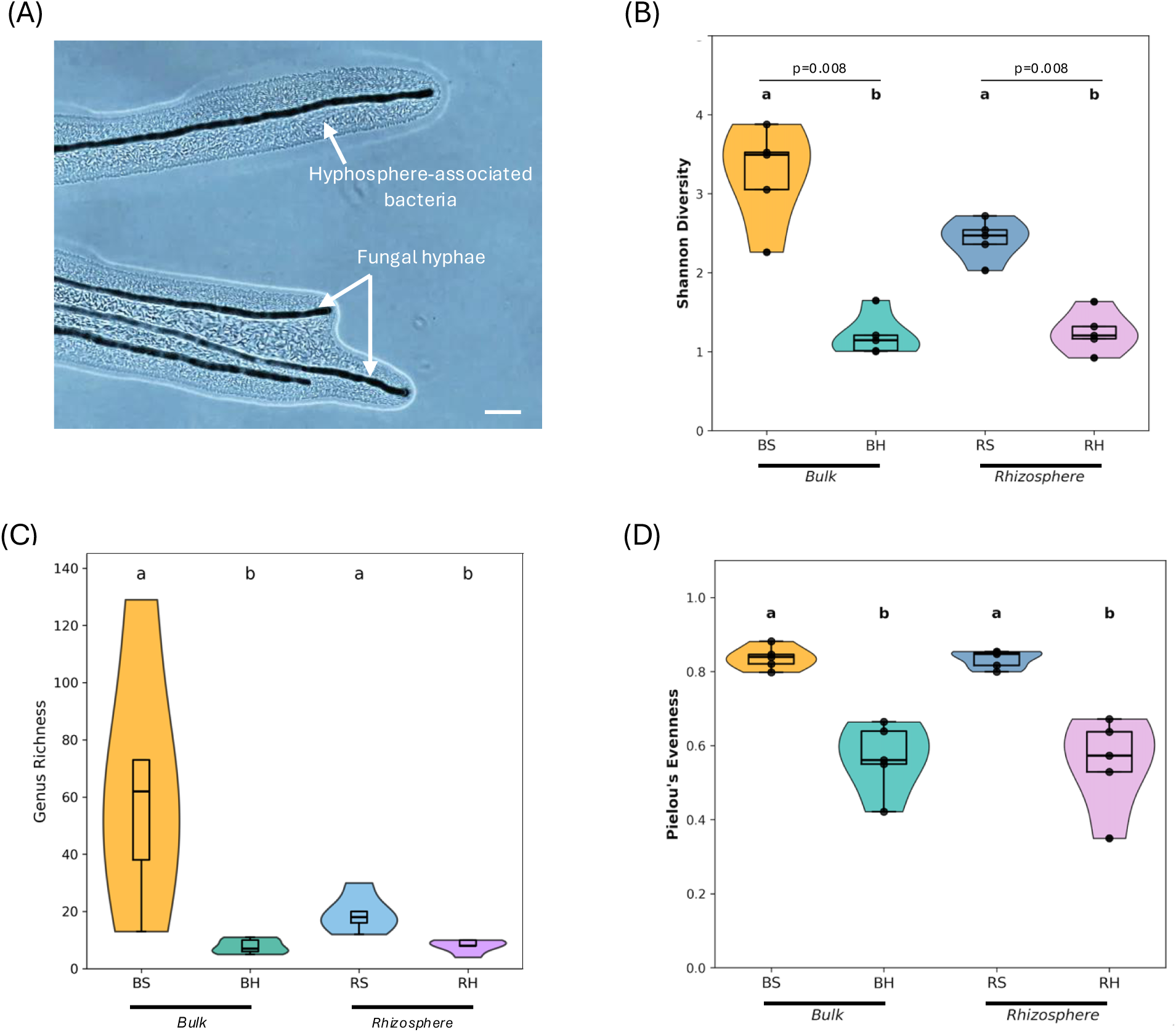
Hyphosphere-associated bacteria and α-diversity patterns in soil and hyphosphere compartments. (A) Brightfield microscopy of FOV4 hyphae obtained from plate cultures initiated from hypha-associated soil microaggregates. Bacterial cells remain attached along the hyphal surface following hyphal outgrowth. Scale bar = 10 μm. (B-D) Alpha diversity metrics of bacterial communities across soil and hyphosphere compartments. Samples include bulk soil (BS), bulk hyphosphere(BH), rhizosphere soil (RS) and rhizosphere hyphosphere (RH), n=5 per group. (B) Shannon diversity was significantly reduced in hyphosphere-associated communities relative to their corresponding soil compartments (Kruskal-Wallis, H = 15.25, p = 0.0016). (C) Genus richness differed among compartments, with highest richness observed in bulk soil and reduced richness in hypha-associated fractions (BS: 63.0 ± 43.5; BH: 7.8 ± 2.6; RS: 19.2 ± 6.7; RH: 8.0 ± 2.4). (D) Pielou’s evenness measured across the same compartments.

### Hyphae recruit a reduced and selective subset of soil bacteria

Hyphosphere communities exhibited a consistent reduction in bacterial diversity relative to surrounding soil compartments. Across both plant-free and plant-associated microcosms, Shannon diversity was significantly lower in hyphosphere samples compared to their respective soil compartments (Kruskal-Wallis H = 15.25, P = 0.0016; Fig. 2B). Pairwise comparisons revealed significant reductions from bulk soil (BS: 3.23 ± 0.45) to bulk hyphosphere (BH: 1.28 ± 0.24, p = 0.008) and from rhizosphere soil (RS: 2.74 ± 0.16) to rhizosphere hyphosphere (RH: 1.45 ± 0.24, p = 0.008). This pattern was accompanied by significantly lower Pielou’s evenness (H = 14.70, p = 0.0021; Fig. 2D), reflecting not only reduced richness but also a more uneven distribution of taxa dominated by a smaller number of enriched genera.

Genus richness analyses further underscored this selective effect. Total genus richness decreased sharply from 169 genera in bulk soil to 22 genera in bulk hyphosphere (87% reduction), and from 38 genera in rhizosphere soil to 29 genera in rhizosphere hyphosphere (24% reduction; Fig 2C). When applying a consistent 0.1% abundance threshold, 93 genera were detected across all compartments (Supplementary Fig. S1), with most detected genera (71%) confined to soil compartments, whereas only 13% were unique to the hyphosphere. Together, these patterns suggest that the hyphal interface constitutes a distinct microhabitat that supports a reduced and selective subset of soil bacteria.

### Soil and hyphosphere compartments exhibit distinct community structures

Beta-diversity analyses visualized as PCoA of Bray-Curtis dissimilarities show compositional differences across compartments BS, BH, RS, and RH communities (Fig. 3A), with significance F = 7.14, R² = 0.572, p = 0.001. Furthermore, we performed pairwise comparisons of all compartments to BS, as it represents the starting inoculum for all the other compartments. These pairwise comparisons show significant compositional divergence between BS and all other compartments (BS vs BH: p = 0.003; BS vs RS: p = 0.008; BS vs RH: p = 0.003), as well as between BH and RS (p = 0.008) and between RS and RH (p = 0.008). Critically, BH and RH did not differ significantly at the whole-community level (R² = 0.128, p = 0.307), indicating convergence of hypha-associated communities despite their different soil origins. Hierarchical clustering and PERMDISP analyses supported the convergence pattern (Supplementary Fig. S2). Hyphosphere samples showed equivalent within-group dispersion (mean distance to centroid: BH = 0.42 ± 0.10, RH = 0.42 ± 0.14), with PERMDISP confirming no significant difference between them (F = 0.063, p = 0.888). This validates that the observed compositional similarity reflects true convergence rather than artifacts of unequal community variability.

**Figure 3.**
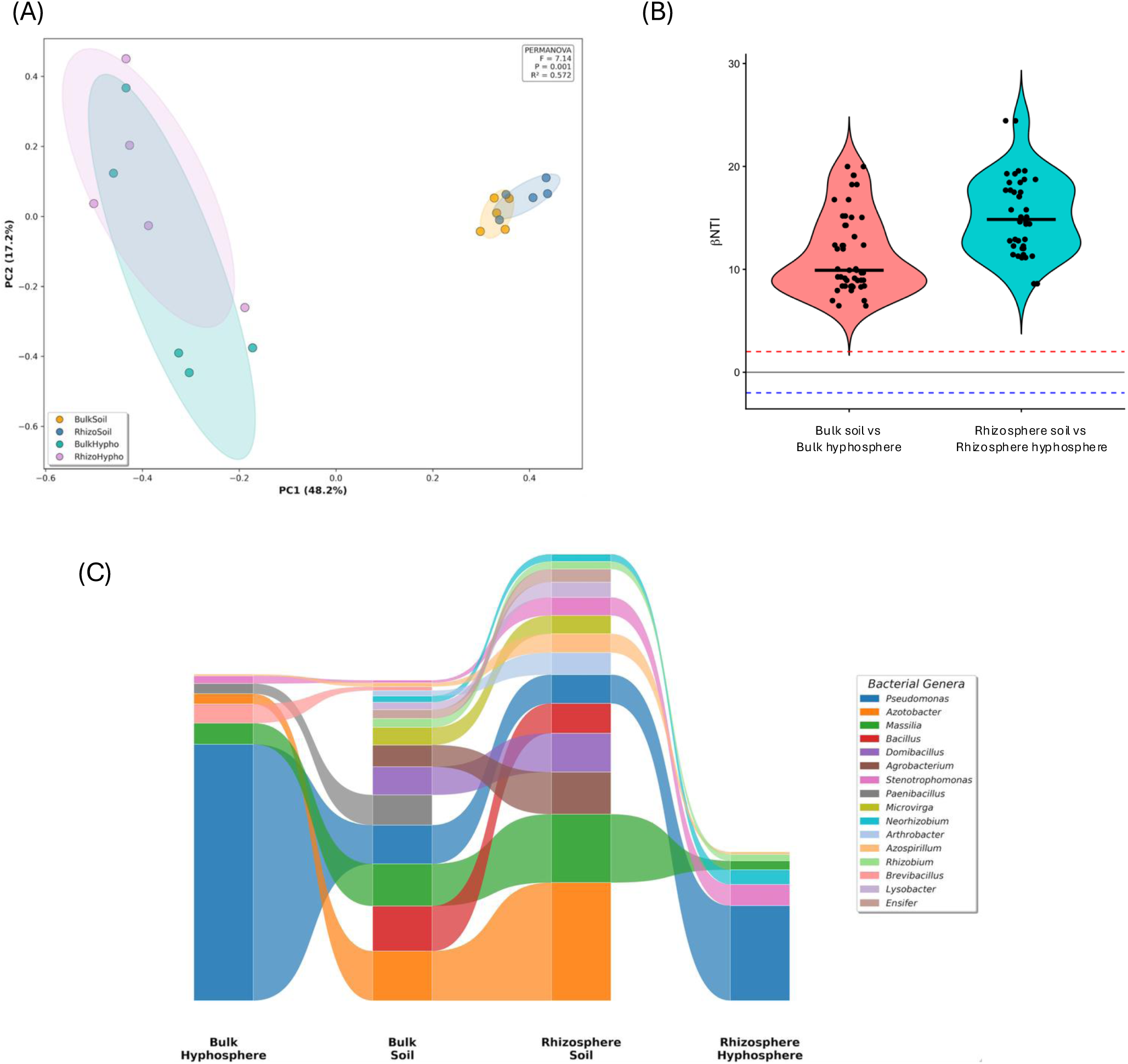
Community structure, phylogenetic turnover and genus-level composition across soil and hyphosphere compartments. (A) Principal coordinates analysis (PCoA) for genus-level bacterial communities from bulk soil (BS), bulk hyphosphere (BH), rhizosphere soil (RS) and rhizosphere hyphosphere (RH) samples based on Bray-Curtis dissimilarity. Points represent individual samples (n = 5 per group) and shaded ellipses indicate 95% confidence regions around group centroids. (B) Beta Nearest Taxon Index (βNTI) values comparing phylogenetic turnover patterns between soil and hyphosphere compartments. Violin plots show the distribution of βNTI values for BS vs BH and RS vs RH transitions. Horizontal dashed lines mark thresholds for homogeneous selection (βNTI < −2) or variable selection (βNTI > +2) relative to a phylogenetic null model. All βNTI comparisons exceeded the +2 threshold, indicating deterministic assembly governed by variable selection during hyphosphere recruitment. (C) Alluvial diagram showing mean relative abundance (%) of dominant bacterial genera across compartments. Flow paths illustrate genus level changes from BS to BH (bulk context) and RS to RH (rhizosphere context). Band width corresponds to mean relative abundance across replicate samples (n = 5 per group). Only genera representing >0.5% mean relative abundance in at least one compartment are shown.

### Phylogenetic turnover patterns during hyphosphere assembly

To evaluate assembly mechanisms underlying hyphosphere formation, we quantified phylogenetic turnover during soil-to-hyphosphere transitions using the beta nearest taxon index (βNTI). Pairwise comparisons revealed positive βNTI values for both bulk soil to bulk hyphosphere transitions (BS vs BH; mean ± SD = 10.93 ± 3.74, median = 9.69, n = 50) and rhizosphere soil to rhizosphere hyphosphere transitions (RS vs RH; mean ± SD = 19.76 ± 11.98, median = 15.96, n = 50; Fig. 3B). All pairwise comparisons (100%) exceeded the +2 threshold for variable selection, with no comparisons falling within the stochastic (−2 to +2) or homogeneous selection (< −2) ranges, indicating that hyphosphere assembly is entirely governed by deterministic processes in both contexts.

The consistently positive βNTI values demonstrate that hyphosphere communities are phylogenetically more dissimilar from their source soil communities than expected under null model predictions, reflecting strong phylogenetic restructuring during bacterial recruitment to fungal hyphae. Notably, RS vs RH transitions exhibited significantly higher βNTI values than BS vs BH transitions (Wilcoxon rank sum test, W = 492, p < 0.001), with nearly twice the mean phylogenetic turnover and substantially greater variability (SD = 11.98 vs 3.74). This elevated and more variable phylogenetic turnover in rhizosphere-associated hyphosphere suggests that plant-conditioned soils generate more heterogeneous assembly outcomes, though both contexts remain dominated by deterministic variable selection processes.

### Hyphosphere communities comprise conserved and compartment-specific bacterial genera

Genus level profiles differed between soil and hyphosphere compartments (Fig. 3C; Supplementary Fig. S3). A subset of bacterial genera, including *Pseudomonas, Enterob*acter, *Achromobacter* and *Pantoea*, was consistently detected at high relative abundance in both bulk and rhizosphere hyphospheres (BH and RH), forming a conserved hyphosphere-associated assemblage (Fig. 4A). Indicator species analysis further supported this pattern, identifying these taxa as reliable markers of hypha-associated microhabitats based on elevated indicator values (IndVal ≥ 0.5, p < 0.05; Fig. 4B). Comparison of genus membership between BH and RH showed substantial overlap of hyphosphere-associated genera (13 shared genera representing ∼50% of total hyphosphere diversity), alongside smaller sets of taxa unique to each compartment (BH-unique: 5 genera and RH-unique: 8 genera; Fig. 4C, left). This indicates that fungal hyphae recruit a partially conserved bacterial consortium across soil contexts, with additional taxa being context dependent, where here it was the presence or absence of the Pima cotton seedling.

**Figure 4.**
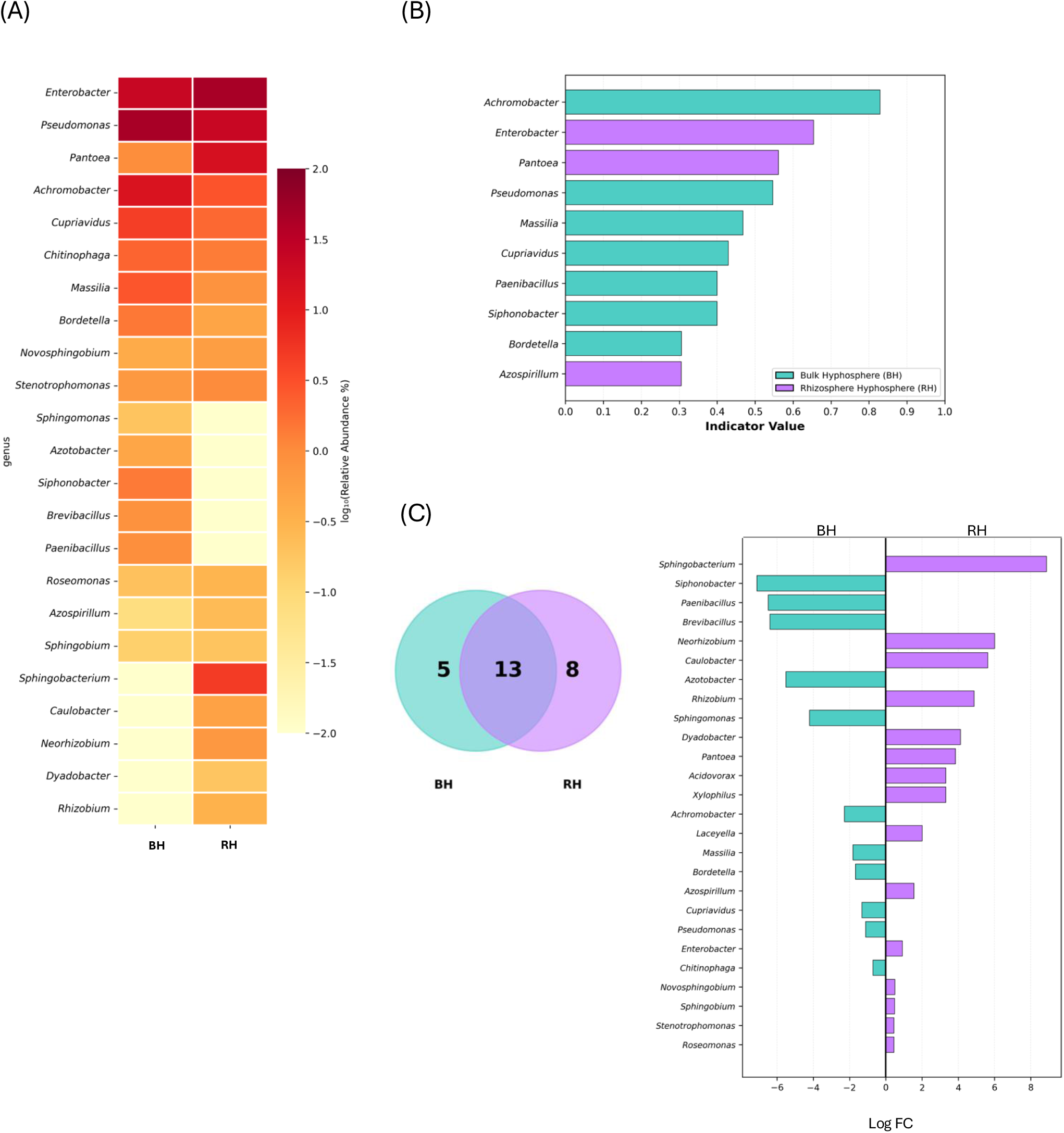
Hyphosphere-resolved genus level composition, indicator taxa and shared vs compartment-specific genera in bulk and rhizosphere hyphospheres. (A) Heatmap showing log₁₀ transformed relative abundances of bacterial genera detected in bulk hyphosphere (BH) and rhizosphere hyphosphere (RH) samples at ≥0.1% relative abundance threshold (n = 23 genera). Values are scaled within each genus to permit comparison of distribution patterns across the two hyphosphere types. (B) Indicator value (IndVal) scores for genera identified by indicator species analysis as characteristic of hyphosphere samples. Bars represent IndVal statistics (range 0-1), reflecting the combined specificity and fidelity of each genus across hyphosphere replicates. (C) Comparison of genus-level membership and relative abundance between BH and RH. Left: Venn diagram showing shared genera and genera unique to each hyphosphere type. Right: Log₂ fold change for genera detected in both compartments, where positive values indicate higher relative abundance in RH and negative values indicate higher relative abundance in BH

Consistent with this pattern, differential abundance analysis demonstrated that several genera were preferentially enriched in BH, including *Siphonobacter* and *Paenibacillus*, whereas others such as *Sphingobacterium* and *Neorhizobium* were relatively more abundant in RH (Fig. 4C, right).

Together, these results show that while hyphal recruitment yields a reproducible core of hyphosphere-associated bacteria, local soil and plant-associated conditions modulate the relative representation of additional taxa within the hyphal microhabitat.

## Discussion

This study demonstrates a novel spatially constrained, soil-mimetic microcosm that can reproducibly recover bacterial communities assembled at the fungal hyphal interface under plant-associated soil conditions. By physically delimiting hyphal outgrowth while preserving soil structure, microbial complexity, and plant influence, the system enabled consistent recovery of hypha-associated microaggregates containing physically attached bacterial cells. Across independent replicates and soil contexts, hyphosphere communities showed reduced diversity, distinct composition relative to surrounding soils, and convergence between bulk and rhizosphere hyphospheres, indicating that fungal hyphae impose a strong and reproducible ecological filter on bacterial community assembly. The microcosm therefore addresses a central methodological limitation in hyphosphere research: the lack of experimental systems that permit spatially resolved sampling of hypha-associated microbiota from intact, plant-conditioned soils.

Our approach addresses methodological limitations through targeted design features that enable spatially constrained recovery under plant-associated soil conditions. The physical barrier provided by cell strainers permits precise isolation of hypha-associated microaggregates, capitalizing on evidence that bacterial communities attached to extraradical fungal hyphae are compositionally distinct from bulk soil communities (Toljander et al., 2006; Scheublin et al., 2010). By recovering microaggregates formed through *in situ* hyphal extension, the system captures bacterial assemblages shaped by authentic soil chemistry, plant-derived gradients, and fungal exometabolite dynamics conditions rarely recapitulated in traditional laboratory systems (Deveau et al., 2018; Jin et al., 2024). This approach complements existing fungal highway and baiting systems that have demonstrated bacterial dispersal and migration processes (Kohlmeier et al., 2005; Banitz et al., 2013; Furuno et al., 2010; Pion et al., 2013; Worrich et al., 2017; Ghodsalavi et al., 2017; Simon et al., 2017). Consistent recovery of FOV4 outgrowth with adherent bacterial cells demonstrates successful capture of authentic hyphosphere communities, with microscopic examination confirming physical bacterial-hyphal associations formed. Unlike column systems relying on indigenous fungi or plant-free baiting approaches, this design allows targeted fungal introduction while preserving soil structure and plant conditioning, supporting both culture-dependent isolation and molecular analyses (Warmink & van Elsas, 2009; Simon et al., 2015).

Community-level analyses showed that bacterial assemblages recovered from hyphosphere compartments were consistently less diverse than those in surrounding soils, indicating that only a subset of the soil microbiota becomes associated with fungal hyphae under the conditions tested. Across both plant-free and plant-associated microcosms, hyphosphere samples exhibited significantly reduced Shannon diversity and Pielou’s evenness relative to their corresponding bulk and rhizosphere soils, reflecting lower richness and increased dominance of a limited number of taxa rather than uniform diversity loss. Comparable contractions in bacterial diversity have been reported in studies of hypha-associated communities in soil and mycorrhizal systems, supporting the view that the hyphosphere represents a more spatially and chemically constrained microhabitat than bulk soil (de Boer et al., 2005; Warmink et al., 2011; Deveau et al., 2018). At micrometer scales, fungal hyphae generate steep gradients in nutrients, water availability, and physicochemical conditions that differ from those experienced by free-living soil bacteria, shaping localized microbial associations at the hyphal interface (Toljander et al., 2006; Tecon & Or, 2017; Nguyen, 2023). Under such constraints, only bacteria capable of persisting at or exploiting the hyphal interface are likely to remain detectable, while many soil taxa may be excluded. The pronounced reduction in pooled genus richness in hyphosphere compartments, together with the predominance of genera detected exclusively in soil samples, supports the interpretation that hyphal interfaces function as selective microbial microhabitats rather than passive collectors of surrounding soil bacteria. Importantly, these patterns were reproducible across independent microcosms and across plant-free and plant-associated conditions, indicating a consistent outcome of hyphal recruitment within the experimental system.

Bacterial community composition differed markedly between soil and hyphosphere compartments, with hypha-associated communities forming discrete assemblages distinct from surrounding soils. Although bulk and rhizosphere soils differed strongly in overall composition, their corresponding hyphosphere communities were more similar to one another, indicating that association with fungal hyphae constrains bacterial community structure and partially reduces differences imposed by initial soil context. Notably, this compositional convergence occurred despite substantially different phylogenetic turnover magnitudes, suggesting that while fungal hyphae impose similar taxonomic filters across contexts, plant-conditioned soils generate more phylogenetically variable assembly outcomes. This compositional convergence suggests functional redundancy at the genus level, where phylogenetically distinct bacterial lineages occupy similar ecological roles on hyphal surfaces. Comparable patterns of reproducible hypha-associated community composition across independent soils and experimental contexts have been reported previously, supporting the view that fungal hyphae can promote similar bacterial assemblages despite divergent starting communities (Emmett et al., 2021; Warmink & van Elsas, 2009; Deveau et al., 2018). Greater dispersion among rhizosphere hyphosphere samples indicates increased variability in recruitment outcomes under plant-conditioned soil conditions, likely reflecting microscale heterogeneity in root-derived inputs, microbial interactions, and resource availability (Toljander et al., 2006; Tecon & Or, 2017; Junier et al., 2021).

Phylogenetic turnover analyses revealed that hyphosphere assembly operates through strong deterministic filtering rather than stochastic processes. The consistently positive βNTI values during soil-to-hyphosphere transitions indicate variable selection, where hyphosphere communities exhibit phylogenetic overdispersion relative to their source soils (Stegen et al., 2012). This pattern suggests that hyphal surfaces impose phylogenetic constraints that favor phylogenetically dissimilar bacterial lineages, potentially reflecting competitive exclusion or niche differentiation among colonizing taxa. The predominance of deterministic over stochastic assembly processes demonstrates that hyphal surfaces represent highly selective microhabitats where phylogenetic structuring, rather than random assembly governs community composition. Taxonomic analyses showed enrichment of specific bacterial lineages exhibiting traits suited to hyphal-associated microenvironments, such as tolerance of localized gradients or affinity for surface attachment, while many abundant soil taxa were excluded from the hyphosphere (de Boer et al., 2005; Warmink et al., 2011; Deveau et al., 2018). These patterns demonstrate that hyphosphere colonization is phylogenetically non-random, with deterministic processes structuring bacterial assemblages at the hyphal interface.

In summary, this microcosm represents a methodological advance in hyphosphere research, providing reproducible, spatially constrained recovery of hypha-associated bacterial communities from intact soil. The system revealed complete deterministic assembly and taxonomic convergence across soil contexts despite phylogenetic divergence. These patterns establish fungal hyphae as reproducible ecological filters that structure bacterial assemblages through phylogenetic overdispersion rather than environmental filtering. By enabling parallel comparisons across soil contexts while maintaining experimental control, this platform facilitates direct testing of whether phylogenetic overdispersion translates to functional niche partitioning, temporal tracking of hyphosphere assembly from colonization and experimental dissection of plant-mediated effects on fungal-bacterial interactions under ecologically realistic soil conditions. We anticipate its simplicity and versatility will support broader adoption of hyphosphere research across agricultural, ecological and pathological systems.

## Author Contributions

GA: validation, formal analysis, methodology, visualization, writing - original draft, review & editing. EN: writing - review & editing. TW: resources, methodology, Writing - review & editing. SP: methodology, writing - review & editing. JS: methodology, writing - review & editing. MB: resources, funding acquisition, writing - review & editing. SAB: conceptualization, validation, resources, funding acquisition, writing - review & editing.

## Acknowledgments

We thank the United States Department of Agriculture (USDA), College Station, Texas, for providing access to growth chamber facilities used in this study.

## Data Availability

All sequencing data generated in this study have been deposited in the NCBI database under BioProject accession number PRJNA1469864.

